# Charge-Altering Releasable Transporters Enable Specific Phenotypic Manipulation of Resting Primary Natural Killer Cells

**DOI:** 10.1101/2020.02.28.970491

**Authors:** Aaron J. Wilk, Nancy L. Benner, Rosemary Vergara, Ole A.W. Haabeth, Ronald Levy, Robert M. Waymouth, Paul A. Wender, Catherine A. Blish

## Abstract

Natural killer (NK) cells are capable of rapid and robust cytotoxicity, making them excellent tools for immunotherapy. However, their recalcitrance to standard transfection techniques has limited both mechanistic studies and clinical applications. Current approaches for NK cell manipulation rely on viral transduction or methods requiring NK cell activation, which can alter NK cell function. Here, we report that non-viral Charge-Altering Releasable Transporters (CARTs) efficiently transfect primary human NK cells with mRNA without relying on NK cell activation. Compared to electroporation, CARTs transfect NK cells two orders of magnitude more efficiently, better preserve cell viability, and cause minimal reconfiguration of NK cell phenotype and function. Finally, we use CARTs to generate highly cytotoxic primary human chimeric antigen receptor NK cells, indicating potential therapeutic utility of this technique. To our knowledge, CARTs represent the first efficacious transfection technique for resting primary NK cells that preserves NK cell phenotype, and can drive new biological discoveries and clinical applications of this understudied lymphocyte subset.

## INTRODUCTION

Natural killer (NK) cells are a group of innate lymphocytes that coordinate and execute the rapid elimination of neoplastic and virus-infected cells^1,2^. Upon activation through a combinatorial array of germline-encoded inhibitory and activating receptors, NK cells can directly kill their targets *via* targeted release of perforin- and granzyme-containing granules, as well as coordinate the downstream immune response by secreting pro-inflammatory cytokines like IFNγ and TNFα^3,4^. The rapid and robust cytotoxicity of NK cells makes them excellent assets for antiviral and anticancer immunotherapy, an enthusiasm most recently bolstered by the use of chimeric antigen receptor (CAR) NK cells in treating CD19^+^ lymphoid cancers with high efficacy and low toxicity^5–13^. Despite their clinical promise, several fundamental facets of human NK cell biology are poorly understood. For example, the molecular underpinnings of human NK cell memory^14–16^ and the precise roles of various NK cell receptors in target cell recognition remain unknown.

NK cells are notoriously difficult to manipulate, presenting challenges both for a deeper understanding of fundamental NK cell biology and for the use of NK cells in clinical applications^17–21^. Most clinical trials involving genetically-modified NK cells utilize viral transduction (NCT02944162, NCT02892695, NCT03056339, NCT03579927)^5,22^. While viral transduction enables stable transgene expression, it is costly, laborious, and induces robust NK cell activation and apoptosis due to triggering of innate nucleic acid sensors, limiting its practicality for mechanistic studies of NK cell biology^8,18,23–28^. Further, in clinical applications, viral transduction carries the risks of sustained adverse effects and oncogenic potential due to insertional mutagenesis, requiring the co-delivery of caspase-based suicide switches^5,6,29^. Most non-viral gene delivery methods, while easier to use for mechanistic studies, have generally only proven efficacious for immortalized NK cell lines with limited efficiency in primary NK cells^30,31,32,33^. The most successful non-viral delivery method for primary NK cells is currently electroporation^8,34^. However, electroporated NK cells generally require cytokine stimulation or expansion on genetically-modified feeder cell lines for adequate transfection efficiency and viability post-electroporation^35–39^, thereby impeding the study of NK biology by altering cellular physiology and phenotype^40^. We therefore sought to develop and optimize an efficacious and bio-orthogonal transfection strategy for primary NK cells.

Recently, we reported the use of Charge-Altering Releasable Transporters (CARTs) for mRNA delivery into T and B lymphocytes^41^. CARTs are multi-block oligomers consisting of a lipid block(s) and a charge-altering block^41–47^. CARTs are initially cationic serving to complex polyanionic nucleic acids but rearrange under basic conditions (pH 7.4) to neutral products, facilitating the release of the anionic cargo^44,47^. Here, we demonstrate that CARTs efficiently transfect primary human NK cells with mRNA without the need for prior cell activation. We use cytometry by time-of-flight (CyTOF) to show that, compared to electroporation, CART-mediated transfection has minimal impact on the NK cell surface repertoire. Further, analysis of NK cell cytotoxicity and cytokine production demonstrates that CART-mediated transfection preserves canonical NK cell functions. Finally, we use this technique to generate robustly cytotoxic human anti-CD19 CAR NK cells, representing a proof-of-concept for this technique’s clinical utility.

## RESULTS

### CARTs transfect resting NK cells more efficiently than commercial reagents

CARTs are synthetic, biodegradable and dynamic delivery vectors, based on di- or tri-block oligomers, consisting of a lipid block(s) followed by a charge-altering block (**Figure 1A**)^41–44^. The charge-altering block is initially cationic for mRNA complexation but rearranges to neutral diketopiperazine small molecules at pH 7.4 facilitating the release of the anionic mRNA while avoiding toxicities associated with persistent cations^41,48,49^. The first generation single-lipid CART D_13_:A_11_ (**1**) has been used for efficient mRNA delivery *in vitro* (>95% transfection in many cultured cells) and *in vivo* (**Figure 1A**)^44^. However, decreased transfection efficiencies were observed in T lymphocytes compared to other cell types, a common trend among transfection reagents^41,50^. Lymphocytes are hypothesized to be difficult for transfection by viral and non-viral vectors in part due to their limited endocytosis and protein translation as well as the detrimental effect of transfection reagents on their viability^51^. However, through high-throughput screening, we previously discovered that CARTs containing both unsaturated C18 oleyl and C9 nonenyl lipids (CART O_5_:N_6_:A_9_ (**2**), CART BDK-O_7_:N_7_:A_13_ (**3**)) resulted in enhanced transfection of T and B lymphocytes, significantly outperforming single-lipid CARTs and commercial reagents^41^. In fact, these mixed-lipid CARTs resulted in >80% transfection of immortalized T lymphocytes *in vitro^41^* as well as ≥1.5% *in vivo* in living mice when administered intravenously^46,52^, significantly outperforming previous transfection technologies for mRNA delivery into T lymphocytes^41,53–55^. We therefore hypothesized that CARTs may represent a novel delivery vector for highly transfection-recalcitrant resting NK cells.

**Figure 1.**
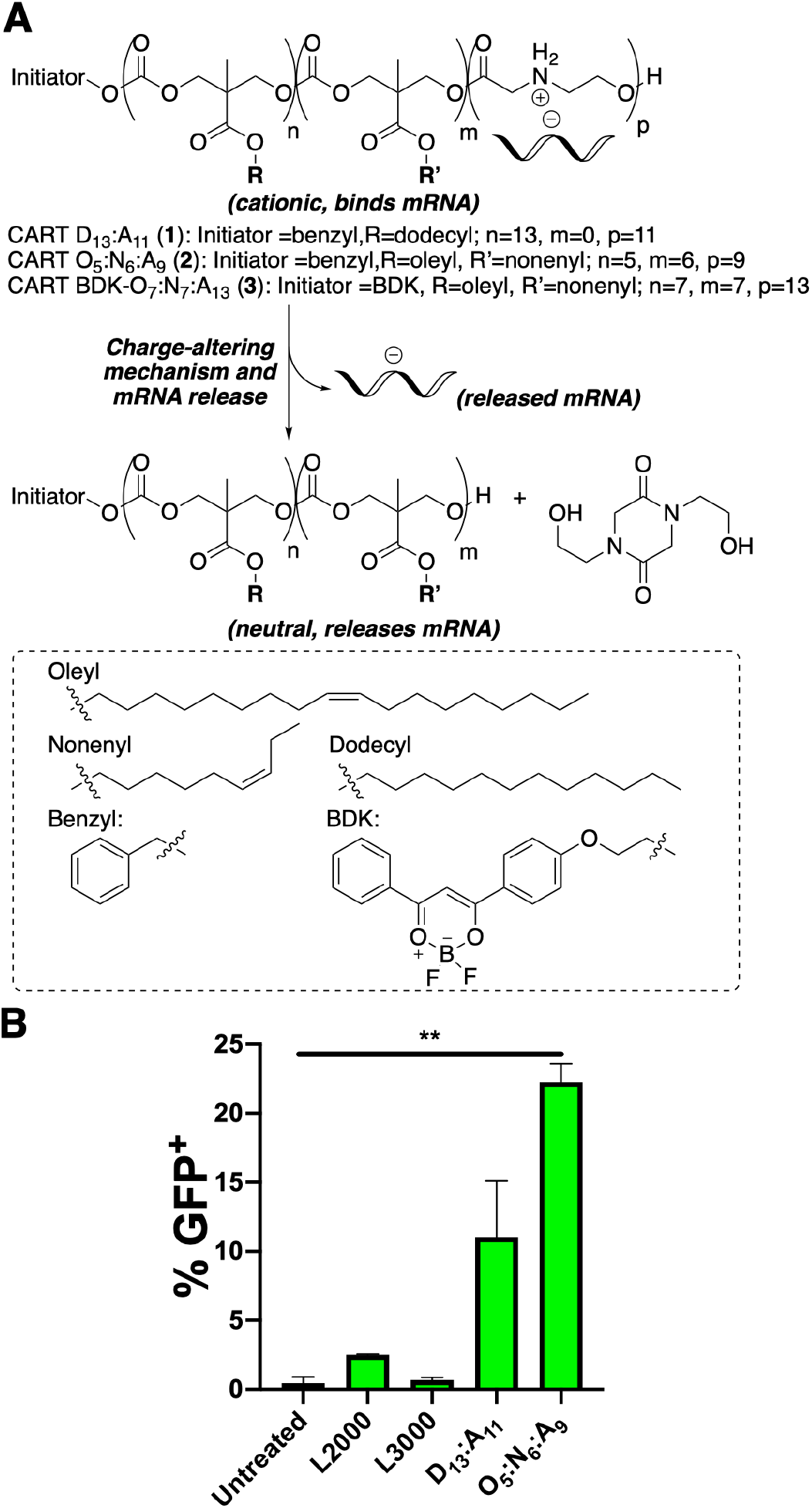
CARTs robustly transfect resting NK cells with mRNA. **A)** Charge-altering rearrangement and structures of single-lipid and mixed-lipid CARTs. **B)** Percent transfection of primary resting human NK cells (100,000 cells/well in 100 μL serum free RPMI) using L2000, L3000, CART D_13_:A_11_ (**1**), or CART O_5_:N_6_:A_9_ (**2**). CART/mRNA polyplexes were formulated at a 10:1 charge ratio (+/−, assuming all ionizable cationic groups are protonated). NK cells were incubated with the complexes for 6 h before analysis by flow cytometry. Error bars represent +/− SD. **, p<0.01 via Welch’s t-test with Bonferroni’s correction for multiple testing. All other comparisons are not significant at the p=0.05 level.

To determine the ability of CARTs to transfect primary human NK cells without activation, we treated isolated NK cells with CART-complexed mRNA encoding a reporter gene. First, we compared the efficacy of single-lipid CART D_13_:A_11_ (**1**) and mixed-lipid CART O_5_:N_6_:A_9_ (**2**) to deliver firefly luciferase (fLuc)-encoding mRNA into primary resting NK cells, where mRNA delivery can be quantified by bioluminescence imaging of the expressed protein in live cells. The CART/mRNA polyplexes were prepared by simple mixing of the CARTs with the mRNA in buffer followed by addition of the polyplexes to NK cells plated in serum-free media. Six hours post-transfection, we found that CART D_13_:A_11_- and O_5_:N_6_:A_9_-transfected NK cells had a ∼20 and ∼50-fold increase in luciferase expression over untreated NK cells, respectively, suggesting efficient cytosolic delivery of the mRNA cargo followed by translation into functional luciferase (**Supplementary Figure 1**).

To more quantitatively compare the ability of CARTs and commercial reagents to deliver reporter mRNA into NK cells, we next turned to green fluorescent protein (GFP) mRNA as both the protein expression and percent of transfected cells could be determined by flow cytometry. We again tested single-lipid CART D_13_:A_11_ (**1**) as well as mixed-lipid CART O_5_:N_6_:A_9_ (**2**) and compared the delivery efficiency of GFP mRNA to commercial lipid nanoparticle (LNP) transfection reagents Lipofectamine 2000 (L2000) and Lipofectamine 3000 (L3000). While minimal GFP was detectable in Lipofectamine-transfected cells (<3%), approximately 10% and 20% of NK cells exhibited GFP expression (GFP^+^) when transfected with CART D_13_:A_11_ (**1**) and O_5_:N_6_:A_9_ (**2**), respectively (**Figure 1B**). This trend is consistent with our previous finding that the mixed-lipid CARTs enhance gene delivery into lymphocytes and indicated that the mixed-lipid CART was the most robust of the transfection methods tested. Therefore we used CART O_5_:N_6_:A_9_ (**2**) and other mixed-lipid CART derivatives for all future experiments.

### CART-mediated transfection of resting NK cells is efficient and scalable

Next, we sought to optimize the efficiency and evaluate the scalability of this technique. We hypothesized that activation and expansion of NK cells would make them more amenable to transfection. Indeed, NK cells activated and expanded for 7 days on the modified K562 feeder line K562-mbIL15-41BBL, a standard method for expansion of primary NK cells^56^, had nearly double the transfection efficacy over resting cells when transfected with CART O_5_:N_6_:A_9_ (**2**)-complexed GFP mRNA (**Supplementary Figure 2C**). To minimize the effect of such expansion and activation protocols on transfected NK cells, we focused our efforts on mRNA delivery to primary resting NK cells for which an efficient transfection method has been lacking. First, we hypothesized that cell density might impact NK cell viability as well as transfection efficiency. We thus examined the impact of NK cell density on viability and transfection efficacy six hours post-transfection with CART O_5_:N_6_:A_9_ (**2**)-complexed GFP mRNA. While we found no significant differences in transfection efficacy when plating between 100,000 and 1,000,000 NK cells per well (1,000,000-10,000,000 cells/mL; **Figure 2A**), we observed that increasing cell density improved the viability of transfected cells (**Figure 2B**). These data suggest that an optimal balance between transfection efficacy and viability might lie at 500,000 NK cells/well, and thus we used this cell density for all following experiments. Additionally, we found that replacing the serum-free media with serum-containing media after 2 or 4 hours did not substantially decrease transfection efficacy while improving cell viability (**Supplementary Figure 2A-B**). This indicates that when high cell viability is critical, shortening the length of the serum-free incubation can preserve viability without sacrificing substantial transfection efficacy. In assays with incubation times longer than 6 hours, we replaced the media with RPMI + 10% FCS after 6 hours.

**Figure 2.**
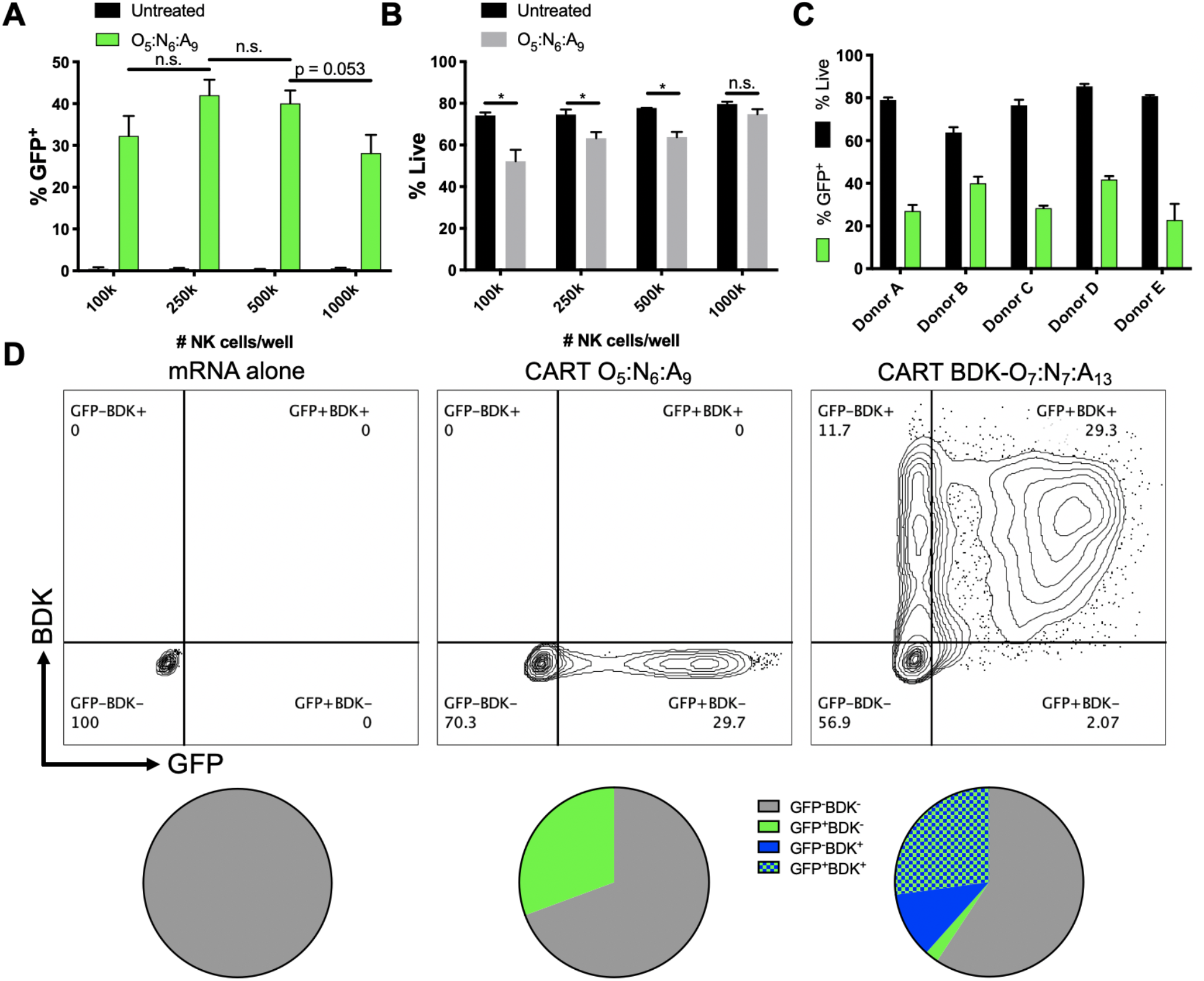
CART transfection of resting NK cells is scalable and selectable. Unless noted, CART/mRNA polyplexes were prepared at a 10:1 (+/−) charge ratio and added to primary resting human NK cells (500,000 cells/well in 100 μL serum free RPMI in 96 well plates) and analyzed by flow cytometry after 6 hours. Effect of NK cell density on percent GFP^+^ cells **(A)** and viability **(B)**. **C)** Effect of NK cell donor on CART O_5_:N_6_:A_9_ (**2**)-mediated GFP mRNA transfection. **D)** Dot plot of GFP expression v. BDK fluorescence when delivering CART BDK-O_7_:N_7_:A_13_ (**3**)/GFP mRNA polyplexes to primary resting NK cells. Percent of NK cells (gated for live cells) that were either BDK^−^GFP^−^, BDK^−^GFP^+^, BDK^+^GFP^−^, or BDK^+^GFP^+^. All measurements were run at least in triplicate. Error bars represent +/− SD. *, p<0.05 by unpaired t-test with Bonferroni’s correction for multiple testing. n.s., not significant at p=0.05.

We next evaluated the extent of inter-donor heterogeneity in cell viability and transfection efficacy using the optimized transfection conditions. In these experiments, we treated primary resting NK cells from five different human donors with CART O_5_:N_6_:A_9_ (**2**)/GFP mRNA polyplexes and tested the transfection efficiency six hours post transfection. Across the five donors tested, the viability ranged between 60-85% and transfection efficacy of GFP mRNA between 20-40% (**Figure 2C**), indicating that CART-mediated transfection of mRNA is likely broadly applicable to human NK cells regardless of donor.

### Co-selection of CART-transfected NK cells

Enrichment for transfected cells often requires co-transfection of a selectable marker that decreases the transfection efficacy of the marker of biological interest^57^. We hypothesized that CARTs may represent a versatile method for co-selection without the delivery of additional cargo due to their chemical tunability. We thus transfected NK cells with GFP mRNA using a mixed-lipid CART (BDK-O_7_:N_7_:A_13_ (**3**)) functionalized with a difluoroboron diketonate (BDK) fluorophore^41,42,52^. BDK is covalently incorporated into every CART oligomer by using it as the alcohol initiator for organocatalytic ring-opening polymerization to provide the initiator-lipid(s)-cationic CART vectors^41^. Additionally, BDK shares many spectral properties with Pacific Blue, enabling it to be easily detected by most flow cytometers. We found that CART BDK-O_7_:N_7_:A_13_ (**3**) had similar transfection efficacy to CART O_5_:N_6_:A_9_ (**2**) in resting NK cells (**Supplementary Figure 2D**), and that the vast majority of GFP^+^ NK cells are also BDK^+^ (**Figure 2D**). These data indicate that CART BDK-O_7_:N_7_:A_13_ (**3**) represents a robust co-selection strategy for transfection of mRNA.

### CART-mediated mRNA transfection causes minimal changes to NK cell proteomic phenotype

NK cells are a remarkably diverse population of innate lymphocytes; previous work has suggested that there may be between 6,000-30,000 subsets of NK cells in a given individual based on the boolean expression of NK cell receptors^58^. This diverse repertoire is stable over time^59^, but can be rapidly and dramatically modulated by a number of common *in vitro* interventions, including low-dose (2 ng/mL) IL-15^40^. We hypothesized that established transfection techniques may alter NK cell proteomic phenotype^60^.

To define how the different transfection techniques impacted the NK cell repertoire, we transfected unactivated NK cells with GFP-encoding mRNA using commercial reagent L2000 or with CART O_5_:N_6_:A_9_ (**2**). We also electroporated primary NK cells that had been stimulated with IL-2 for 24 hours to match previously published protocols^35^. We found a large discrepancy between the dose of mRNA we used for CART-mediated transfection (31 ng/well) and the dose published for electroporation (10,000 ng/well)^34^, and thus IL-2 stimulated NK cells were electroporated with both doses of mRNA. Additionally, for the most accurate representation of the baseline NK cell repertoire, we included as a negative control NK cells maintained in RPMI with 10% FCS. Six hours post-transfection, we analyzed cell viability and transfection efficacy by flow cytometry and stained for CyTOF to evaluate NK cell phenotype.

Between the tested transfection techniques, GFP expression was only detected in the CART-transfected NK cells and the NK cells electroporated with the 322-fold higher dose of mRNA, indicating the CART-mediated transfection is several orders of magnitude more efficient in terms of reagent use (**Supplementary Figure 4A-B**). We observed uniform decreases in expression of most NK cell receptors in the three conditions that were incubated in serum-free media (untreated cells in serum-free media, L2000-, and CART-transfected cells; **Supplementary Figure 4C**), consistent with serum starvation. There were no clear phenotypic differences between CART-transfected cells and NK cells maintained in serum-free media (**Supplementary Figure 4C**), indicating that CARTs likely do not cause unwanted off-targets effects on NK cell phenotype. In contrast, electroporated cells had undergone substantial proteomic reconfiguration, losing expression of five killer cell immunoglobulin-like receptors (KIRs), three natural cytotoxicity receptors (NCRs), as well as markers of NK cell maturity (NKG2C and CD57). Electroporated cells also upregulated activation markers CD38 and CD69, inhibitory receptor TIGIT, activating receptor NKG2D, and target cell apoptosis-inducing Fas-L (**Supplementary Figure 4C**).

As it appeared likely that the changes in NK cell phenotype induced by L2000- and CART-mediated transfection were due to serum starvation, we hypothesized that resting the NK cells in serum-containing media may allow them to regain their baseline phenotype. To test this, we transfected isolated NK cells with GFP-encoding mRNA *via* L2000, electroporation, or CART O_5_:N_6_:A_9_ (**2**) as above, changed the media in all conditions 6 hours post-transfection, let the cells rest for 12 hours, and then performed flow cytometric analysis for viability and transfection efficacy, and CyTOF for NK cell repertoire analysis. At 18 hours post-transfection, we observed that CART-transfected cells maintained substantially higher viability than electroporated cells (**Figure 3A**), and that CART-transfected cells and cells electroporated with the higher dose of mRNA had comparable transfection efficacies (**Figure 3B**). Non-linear dimensionality reduction indicated that CART-transfected NK cells were phenotypically similar to untreated NK cells, suggestive of minimal off-target effects (**Figure 3C**). However, electroporated cells were dramatically phenotypically distinct from the remaining conditions (**Figure 3C**). Hierarchical clustering by mean marker expression did not separate CART-transfected NK cells from untreated serum-replete, untreated serum-starved, or lipofectamine-treated cells, while the first branch of this clustering separated all electroporated samples (**Figure 3D**). There were no clear and consistent changes in mean marker expression in CART-transfected NK cells. Electroporated cells across all donors, on the other hand, displayed reduced expression of activating NK cell receptors 2B4, NKp46, and NTB-A, and increased expression of activation markers CD38, CD69, and HLA-DR, as well as Fas-L and the signaling adaptor FcεRIγ (**Figure 3D**).

**Figure 3.**
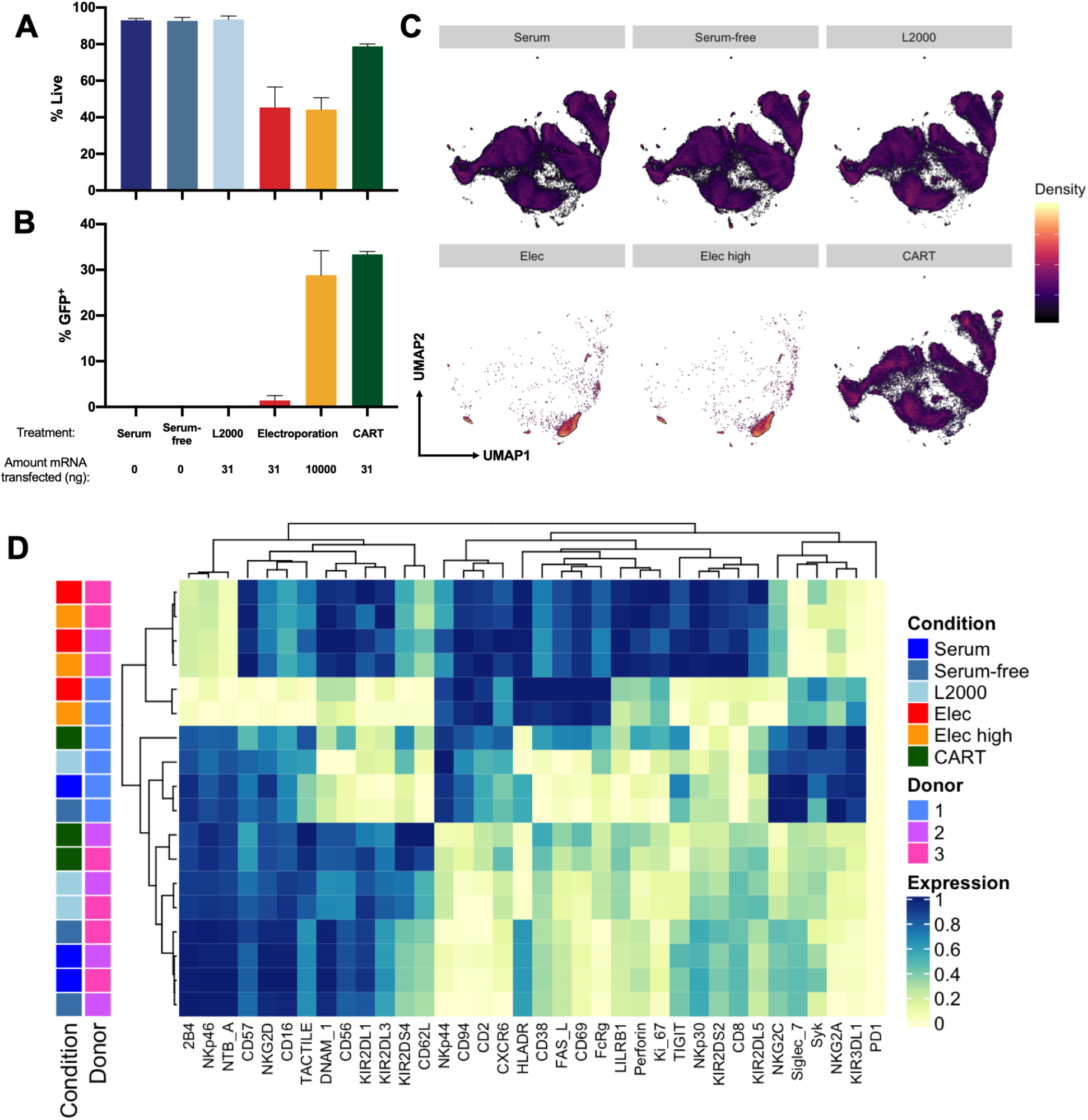
CART transfection outperforms electroporation and causes minimal reconfiguration of NK cell phenotype. NK cells (500,000 cells per well) were transfected with GFP-encoding mRNA *via* L2000 (31 ng mRNA/well), electroporation (31 ng/well (Elec) or 10,0000 ng mRNA/well (Elec high)), or CART (**2**) (31 ng mRNA/well). Flow cytometric analysis of NK cell viability **(A)** and transfection efficacy **(B)** 18 hours post-transfection with GFP-encoding mRNA. **C)** UMAP dimensionality reduction of mass cytometry data faceted by treatment condition. **D)** Heatmap of mean marker expression, with samples and markers hierarchically clustered.

To analyze these changes in NK cell repertoire with more granularity, we clustered the cells using the Louvain method for community detection, identifying 22 clusters in the dataset (**Figure 4A**). We found that all clusters present in untreated NK cells were also represented in CART-transfected NK cells (**Figure 4D**), and that CART-transfected NK cells congruently have a comparable Simpson diversity index to untreated cells (**Supplementary Figure 5**). However, >50% of cells in the electroporated conditions belonged to one of only 3 clusters (3, 21, and 22; **Figure 4D**), reflected in a substantially lower Simpson index than for untreated cells (**Supplementary Figure 5**). The three clusters enriched in the electroporated samples have in common the highest levels of CD69 expression in the dataset. Cluster 21 represents a group of CD56^dim^ NK cells that are separated from its next nearest cluster by upregulation of HLA-DR, NKp30, CD94, TIGIT, Fas-L, CD38, and Perforin. Cluster 22 is a cluster of CD56^bright^ NK cells that, compared to the other CD56^bright^ NK cells in the dataset, notably express lower levels of the homing receptor CD62L (**Figure 4B-C**). Collectively, these data indicate that electroporation dramatically reconfigures the NK cell repertoire. CART-mediated transfection, on the other hand, appears to be a more bio-orthogonal technique that does not cause off-target phenotypic reconfiguration of NK cells while yielding vastly superior transfection efficacy in comparison with lipofectamine. Thus, CART-mediated transfection can be used to manipulate NK cell phenotype efficiently while limiting off-target effects.

**Figure 4.**
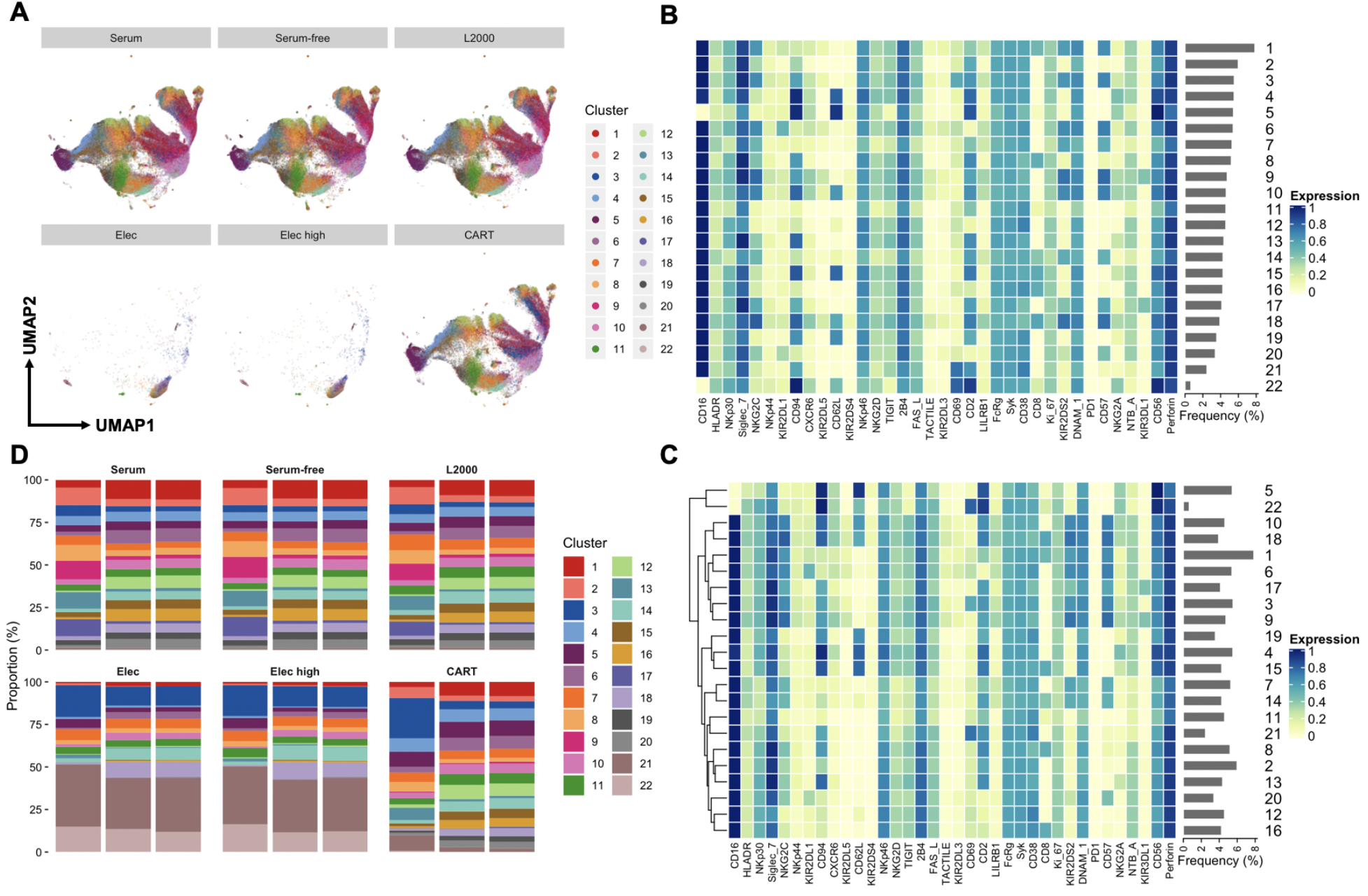
Louvain clustering of mass cytometry data reveals that CART-transfected NK cells maintain NK cell repertoire diversity. NK cells (500,000 cells per well) were transfected with GFP-encoding mRNA *via* L2000 (31 ng mRNA/well), electroporation (31 ng/well (Elec) or 10,0000 ng mRNA/well (Elec high)), or CART (**2**) (31 ng mRNA/well). **A)** UMAP dimensionality reduction plot colored by identified clusters. Heatmap of mean marker expression per cluster, with rows ordered by cluster number **(B)** or with rows clustered hierarchically **(C)**. **D)** Proportions of cells belonging to each cluster.

**Figure 5.**
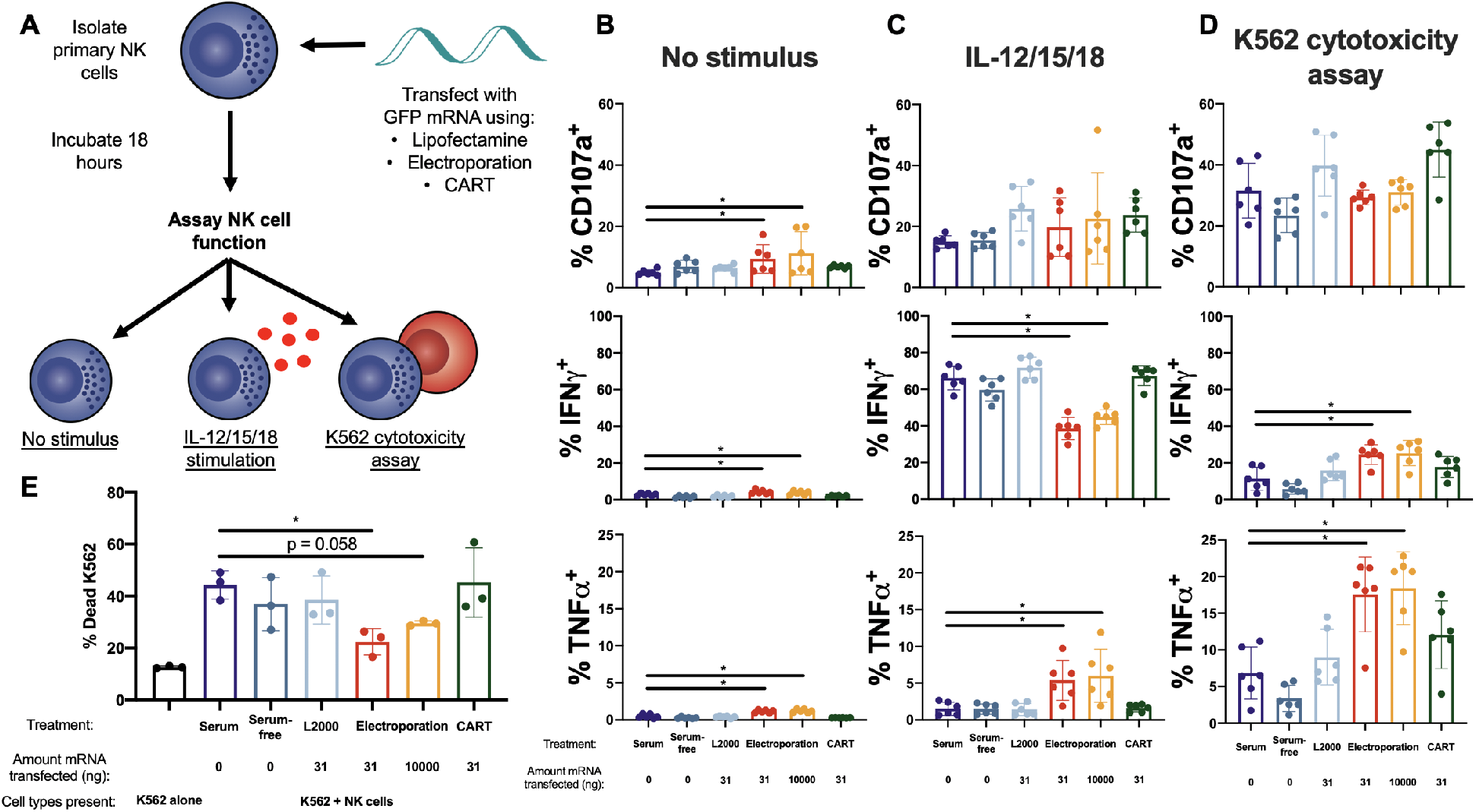
NK cell cytotoxic potential and cytokine production profiles are preserved after CART transfection. **A)** Isolated NK cells (500,000 cells per well) were transfected with GFP-encoding mRNA *via* L2000 (31 ng mRNA/well), electroporation (31 or 10,000 ng mRNA/well), or CARTs (31 ng mRNA/well), and incubated for a total of 18 hours before assay of functional capacity by IL−12, −15, and −18 cytokine stimulation and K562 cytotoxicity assay. Percentage of NK cells with surface CD107a, intracellular IFNγ, or intracellular TNFα without stimulus **(B)**, after IL−12, −15, and −18 cytokine stimulation **(C)**, or after K562 cytotoxicity assay at an effector:target (E:T) ratio of 1:3 **(D)**. *, p<0.05 via Wilcoxon matched pairs signed-rank test. **E)** Percentage of dead K562 cells after 10:1 E:T co-culture with transfected NK cells. *, p<0.05 via paired t-test.

### CART transfection preserves NK cell functional phenotype

We next sought to determine how Lipofectamine-, electroporation-, and CART-mediated transfection of mRNA impacted the ability of NK cells to perform their canonical functions. We therefore transfected resting or IL-2 stimulated primary NK cells with GFP-encoding mRNA using L2000, CART O_5_:N_6_:A_9_ (**2**), or electroporation as described above. We then changed the media 6 hours post-transfection and let the cells rest for another 12 hours before assaying two classical aspects of NK cell function: cytotoxicity in response to co-culture with K562 cells and cytokine production in response to IL-12, −15, and −18 stimulation (**Figure 5A**).

Across all experimental conditions, CART-transfected resting NK cells are indistinguishable from their untransfected counterparts (**Figure 5B-D**). CART-transfected NK cells respond to IL−12, −15, and −18 stimulation with the same magnitude of IFNγ production as untreated NK cells (**Figure 5C**). In response to co-culture with K562 cells, CART-transfected NK cells degranulate, produce IFNγ and TNFα, and kill the targets comparably to untreated NK cells (**Figure 5D-E**). This demonstrates that the CART transfection protocol does not alter canonical NK cell functions.

Electroporated NK cells, in contrast, were functionally distinct from untreated cells. In NK cells that received no further stimulus (not treated with IL−12, −15, −18 or co-cultured with K562 cells), electroporated NK cells expressed higher baseline levels of CD107a, IFNγ, and TNFα (**Figure 5B**), indicating spontaneous degranulation and pro-inflammatory cytokine production. Further, electroporated NK cells expressed significantly less IFNγ and more TNFα in response to IL-12, −15, and −18 stimulation (**Figure 5C**), suggesting a skewing of NK cell cytokine response. Electroporated cells also expressed significantly higher levels of both IFNγ and TNFα compared to untreated cells when co-cultured with K562 cells (**Figure 5D**). Despite this, electroporated cells were slightly less able to kill their K562 targets (**Figure 5E**). These data collectively indicate that, unlike electroporation, CART-mediated transfection preserves canonical NK cell functions, making it a particularly suitable technique for the study of NK cell biology.

### Generation of highly cytotoxic CAR NK cells using CARTs

CAR NK cells have shown promise in several clinical trials^8,9^. We hypothesized that CARTs could be used to generate human CAR NK cells *in vitro*, and designed and synthesized an mRNA encoding the anti-CD19-41BB-CD3ζ CAR^61^. We transfected isolated resting human NK cells with this CAR-encoding mRNA using L2000 or CART BDK-O_7_:N_7_:A_13_ (**3**), in order to track transfected NK cells for downstream analysis. We then rested the cells for a total of 18 hours before a 6 hour co-culture with the target cells of interest.

We found that CART-mediated transfection of the anti-human CD19 CAR mRNA (hCAR) induced robust expression of the CAR on the NK cell surface (**Figure 6A**). With an average of 12% of CART-transfected cells expressing the CAR (**Figure 6A**), we detected a significant increase in the percentage of dead CD19^+^ Raji target cells (a human B lymphoblast cell line) when co-cultured with CAR mRNA CART-transfected NK cells (**Figure 6B**). This demonstrates that CARTs can be used to generate cytotoxic human CAR NK cells.

**Figure 6.**
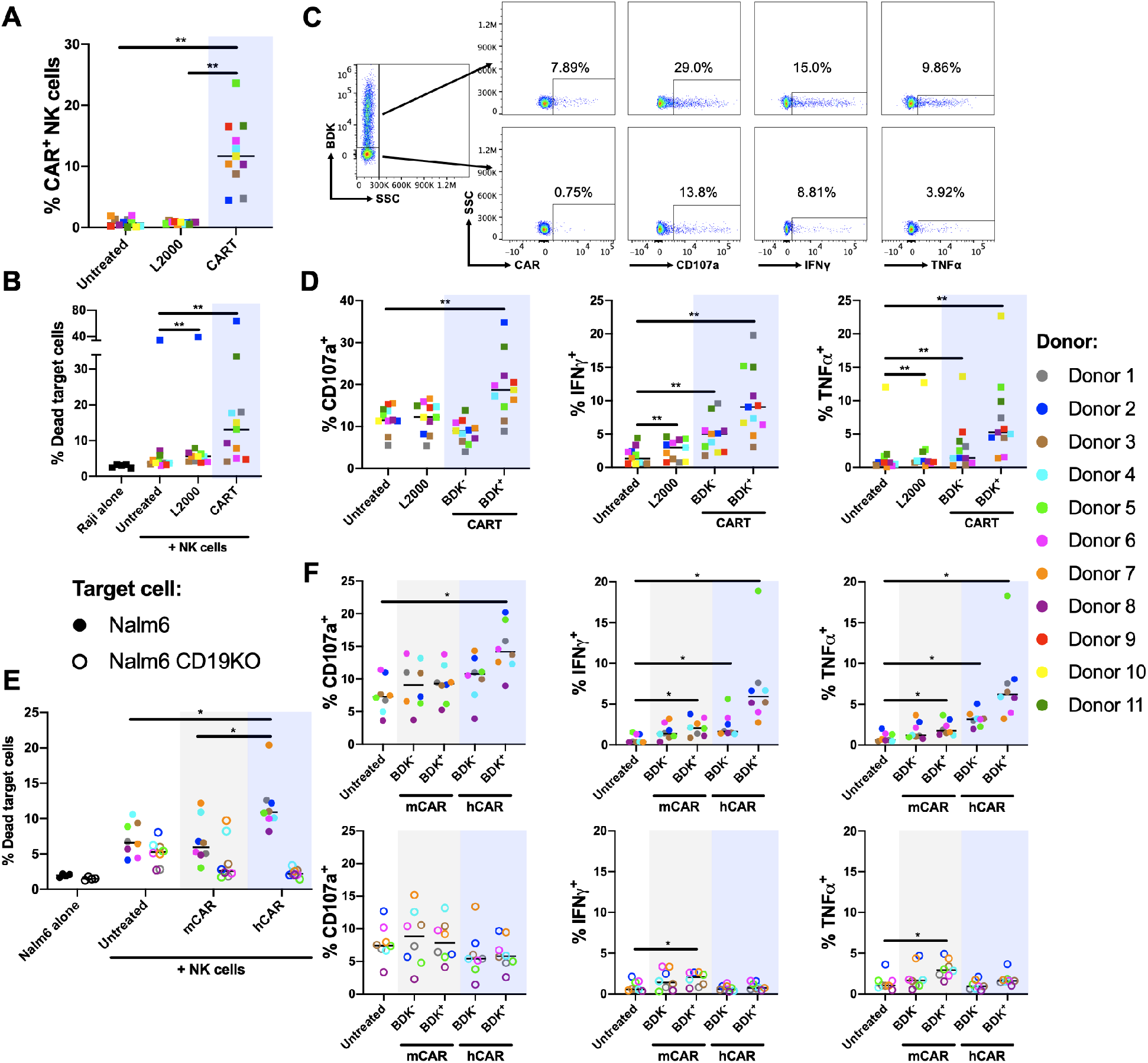
CART-mediated transfection of anti-CD19 CAR generates cytotoxic human CAR NK cells. Isolated primary resting human NK cells were transfected with an mRNA encoding a anti-human CD19-41BB-CD3ζ CAR (hCAR). The cells were incubated for a total of 18 hours before being co-cultured with Raji (■), wild-type Nalm6 (●), or CD19KO Nalm6 (○) target cells for 6 hours followed by flow cytometric analysis. **A)** anti-human CD19 CAR (hCAR) expression by NK cells in the absence of target cells, as detected by FITC-conjugated soluble human CD19. **B)** Percentage of dead Raji cells after E:T 10:1 co-culture with transfected NK cells. **C)** Representative flow cytometry plots of CD107a, IFNγ, and TNFα between BDK^−^ and BDK^+^ populations of CART-transfected NK cells co-cultured with Raji cells at E:T 1:3. This analysis is quantified across all n = 11 donors in **(D)**. **E)** Percentage of dead wild-type or CD19KO Nalm6 cells after E:T 20:1 co-culture with transfected NK cells. **F)** CD107a, IFNγ, and TNFα expression after E:T 1:3 co-culture with Nalm6 cells. For E) and F), n = 8. For all panels, *, p<0.05; **, p<0.01 by Wilcoxon matched-pairs signed rank test with Bonferroni correction for multiple testing.

Although the percentage of BDK^+^ NK cells was consistent when cultured in the presence or absence of Raji target cells, co-culture with the target cells induced a significant decrease in the percentage of NK cells expressing the CAR on the surface (**Supplementary Figure 6**). This suggests that the surface-expressed CAR is likely being ligated in the presence of its cognate antigen and undergoing endocytic internalization and recycling, a process that has been described for several NK cell receptors upon ligation^62–64^. Thus, to investigate the impact of CAR expression on NK cell degranulation and cytokine production in response to Raji cells, we compared CD107a, IFNγ, and TNFα expression by BDK^−^ vs. BDK^+^ CART-transfected NK cells (**Figure 6C**). There was significantly higher degranulation amongst BDK^+^ CART-transfected NK cells compared to untreated cells (**Figure 6D**), suggesting a higher frequency of potentially cytolytic interactions among CAR^+^ NK cells. While BDK^+^ CART-transfected NK cells also expressed the highest levels of IFNγ and TNFα, the presence of CAR NK cells appeared to induce bystander cytokine secretion, as BDK^−^ NK cells from the CART-transfected condition expressed significantly more IFNγ and TNFα than at baseline (**Figure 6D**).

To confirm that this enhanced cytotoxicity and cytokine production was due to ligand-specific recognition by the transfected CAR, we tested the ability of NK cells transfected with anti-murine CD19 CAR (mCAR) to recognize human CD19^+^ target cells (Nalm6), and we tested the ability of CAR-transfected NK cells to respond to human CD19^−^ target cells (CD19KO Nalm6). We found that only anti-human CD19 CAR (hCAR)-transfected NK cells exhibited enhanced killing of human CD19^+^ Nalm6 target cells; murine anti-CD19 CAR (mCAR) did not exhibit enhanced killing of human CD19^+^ Nalm6 cells (**Figure 6E**). In addition, human anti-CD19 CAR (hCAR)-transfected NK cells were significantly more lethal to human CD19^+^ Nalm6 cells relative to CD19^−^ target cells (Nalm6 CD19KO, **Figure 6E**). Furthermore, only hCAR-transfected NK cells had substantially enhanced levels of degranulation and IFNγ and TNFα production, and this trend was observed only after co-culture with CD19^+^ Nalm6 target cells (**Figure 6F**). There was a modest but significant increase in IFNγ and TNFα production by BDK^+^ mCAR-transfected NK cells, which we attribute to tonic signaling of the transfected CAR as it is present both in response to parental and CD19KO Nalm6 target cells (**Figure 6F**). However, mCAR-transfected NK cells displayed no increase in degranulation or target cell killing (**Figure 6E-F**), implying that tonic CAR signaling does not impact cytolytic mechanisms. Collectively, these studies demonstrate that CARTs can be used to generate CAR NK cells with sharpened effector functions.

## DISCUSSION

Here we describe a new technique for mRNA delivery into difficult-to-transfect primary NK cells. CART-mediated transfection is more than 300 times more efficient and better preserves cell viability compared to electroporation, the most commonly used non-viral mRNA delivery technique. We have extensively characterized the phenotype of CART-transfected NK cells by high-dimensional mass cytometry as well as multiple assays of canonical NK cell functions. Using these methods we show that CART-mediated transfection does not induce off-target effects and thus represents a technique for the specific phenotypic manipulation of NK cells. To our knowledge, this work represents the first demonstration of a non-viral primary NK cell transfection that does not require prior cell activation, and thus may currently be the only method for bio-orthogonal manipulation of primary NK cells.

As electroporation is the most widely used non-viral transfection method for primary lymphocytes, we thoroughly benchmarked CART transfection of NK cells with electroporation. Our data revealed that electroporation may induce numerous off-target effects on the NK cell repertoire, skew NK cell cytokine production profiles, and deleteriously impact NK cell killing capacity. In these studies, we pre-treated the electroporated NK cells with IL-2, as NK cell activation appears to be required for post-electroporation viability^35–39^. Interestingly, the phenotypic changes induced by electroporation were not congruent with the alterations expected to be induced by IL-2 treatment alone. For example, IL-2 treatment results in upregulation of NKp30 and NKp44^40^, but the expression of these receptors was lost 6 hours post-electroporation (**Supplementary Figure 4C**). Additionally, IL-2 treatment does not cause spontaneous secretion of IFNγ^40^ as we observed in electroporated NK cells (**Figure 5B**). Ultimately, these data point to electroporation-specific off-target effects on NK cell phenotype, making it unsuitable for rigorously answering open questions of NK cell biology.

A bio-orthogonal transfection strategy for resting human NK cells is necessary to begin addressing fundamental mechanistic questions in human NK cell biology. First, the contributions of many individual NK cell receptors in the recognition of specific target cells remain poorly understood. For example, the activating NK cell receptor NKp46 is reported to recognize influenza hemagglutinin^65,66^, but mouse models of NKp46 deficiency have yielded differing phenotypes upon influenza infection^67,68^. Similar conundrums exist with other NK cell receptors, including activating KIR3DS1 and inhibitory KIR3DL1, which despite having nearly identical extracellular domains and opposing downstream signaling, are both strongly associated with slower progression to AIDS in HIV infection^69,70^. Another situation in which mechanistic studies enabled by bio-orthogonal transfection techniques could enhance our understanding of NK cell biology is in the setting of human NK cell memory. Populations of human NK cells with memory-like phenotypes are deficient in specific transcription factors and signaling adaptors^14,15^, but whether these deficiencies are necessary or sufficient for memory-like functionality remains unclear. As these questions involve molecules and phenotypes that are rapidly modulated by common biological perturbations like cytokine treatment^40^, it is important to use a transfection technique that maintains native NK cell physiology. Our high-dimensional analysis of NK cell phenotype and function after CART-mediated transfection indicates that CARTs do not induce unwanted off-target effects, ideally suiting CARTs to begin addressing basic questions in NK cell biology.

While the ability to transfect primary resting NK cells represents a significant advance for understanding human NK cell biology, CART-mediated transfection may also enhance clinical efforts. We demonstrated the potential clinical utility of CARTs by generating anti-CD19 human CAR NK cells *in vitro*. These CAR NK cells are potently cytotoxic and are more robustly activated in response to CD19^+^ target cells than their untransfected counterparts. As the transfection is mRNA-mediated and thus transient, this approach might mitigate the need for a suicide switch, as was used in the recent clinical trial of virally transduced CAR NK cells^5^. Further studies will need to examine whether sufficient numbers of CART-generated CAR NK cells can be readily generated, and to assess their lifespan *in vivo* in order to determine if this technique would be superior to lentiviral transduction for CAR NK cell preparation. As we have previously demonstrated that CARTs can be used to deliver mRNA *in vivo* with a high specificity for mRNA expression in the spleen^41,46,52^, *in vivo* CART-mediated delivery of CAR-encoding mRNA may provide an alternative therapeutic approach for the generation of CAR NK cells as a strategy for personalized cancer immunotherapy.

Finally, an attractive aspect of the CART technology is that CARTs can be readily synthesized on scale in two steps from monomers and their lipid and charge composition can be readily varied as dictated by research and clinical needs. The conversion of the CART oligomers to nanoparticle complexes upon simple mixing with mRNA, the efficacy and selectivity of delivery, accommodation of a range of nucleic acid cargos including CRISPR, and lack of acute immunogenicity offer further potential advantages for their use^41,42,45,52^.

In conclusion, we report the use of CARTs to phenotypically manipulate primary human NK cells *in vitro* with a minimal effect on native NK cell biology. This method shows enhanced transfection and improved cell viability compared to other non-viral transfection methods (electroporation, Lipofectamine) using the same dose of mRNA. We then applied the developed CART technology to generate CAR NK cells, which exhibit enhanced killing of the target cells. We are continuing to develop new CARTs with varying architectures and targeting ligands to further enhance gene delivery into primary cells and enable further biomedical research and therapeutic use.

## MATERIALS AND METHODS

### Isolation and culture of NK cells and cell lines

Peripheral blood mononuclear cells (PBMCs) were isolated by Ficoll-Paque (GE Healthcare) density gradient centrifugation and cryopreserved in 90% FBS + 10% DMSO (v/v). PBMCs were thawed at 37°C in complete RPMI-1640 media (supplemented with 10% FBS, L-glutamine, and Penicillin-Streptomycin-Amphotericin; RP10) containing benzonase (EMD Millipore). NK cells were purified by magnetic bead isolation via negative selection according to the manufacturer’s specifications (Miltenyi, cat. 130-092-657). Unless otherwise noted, NK cells were maintained in complete RP10 media without any additional cytokines to ensure a resting state.

K562 cells, Raji cells (ATCC), wild-type Nalm6 cells, and CD19KO Nalm6 cells (a gift from the lab of Prof. Crystal Mackall) were used from passage 2-10 and maintained in RP10. To confirm deletion of CD19 in CD19KO Nalm6 cells, this cell line was stained with anti-human CD19-PerCP-Cy5.5 (BioLegend, Clone HIB19). All cell culture was performed at 37°C/5% CO_2_ in a humidified environment.

### Synthesis of CARTs and preparation of CART/mRNA polyplexes for cellular assays

CART O_5_:N_6_:A_9_ (referred to as O_5_-*b*-N_6_:A_9_ in a previous report^41^), CART BDK-O_7_:N_7_:A_13_, and CART D_13_:A_11_ were prepared as previously described^41,44^. The CARTs were stored as 2 mM stock solutions in DMSO at −20 °C. As a sample procedure to prepare the CART/mRNA polyplexes, we first diluted 0.42 μL of the GFP mRNA (Trilink, stored at 1 μg/μl) into 7.26 μL of PBS (pH 5.5) in a 0.6 mL eppendorf tube. The PBS pH 5.5 was prepared by adjusting the pH of RNAse free PBS 1X (pH 7.4) to pH 5.5 with HCl. To the solution of mRNA was added 0.72 μL of CART O_5_:N_6_:A_9_ (stored as a 2 mM stock solution in DMSO) to achieve a charge ratio of 10:1 (+/−, assuming all ionizable cationic groups are protonated). After mixing by finger vortex for 15 seconds, the CART/mRNA polyplexes were added to the cells of interest (typically 0.625 μL was added to achieve 31 ng of mRNA per well). Following addition of CART/mRNA polyplexes to the wells, the NK cells were incubated for 6 hours in serum-free media.

### Lipofection

Lipofectamine 2000 and Lipofectamine 3000 were purchased from ThermoFisher and used according to manufacturer’s instructions. The same final dose of mRNA per well was used for the Lipofectamine conditions as the CART conditions in all comparative assays.

### Electroporation

Primary NK cells were electroporated as previously described^35^. Briefly, isolated NK cells were cultured in 1000 IU/mL human IL-2 (R&D Systems) for 24 hours. NK cells were then centrifuged, resuspended in freshly-made electroporation buffer (5 mM KCl, 15 mM MgCl_2_, 15 mM HEPES, 150 mM Na_2_HPO_4_/NaH_2_PO_4_, 50 mM mannitol, pH 7.2), mixed with the appropriate volume of mRNA, and transferred into a 100 μL nucleofector cuvette at a final density of 5,000,000 NK cells/mL. Cells were then electroporated with the 4D Nucleofector System (Lonza) using pulse code CM-137 and buffer code P3. After electroporation, 100 μL of pre-warmed complete media was added to each cuvette, incubated at 37 °C/5% CO_2_ for 10 minutes, and transferred to a 96-well plate for further culture.

### mRNA synthesis

eGFP mRNA (996 nucleotides) was purchased from TriLink and used without further purification.

CAR-encoding mRNA was transcribed from linearized plasmids encoding anti-human CD19 CAR^61^ and anti-mouse CD19 CAR^71^ using T7 RNA polymerase (HiScribe™ T7 High Yield RNA Synthesis Kit, New England Biolabs). The mRNAs were transcribed to contain the 5′ UTR derived from the human a-globin 5′ leader RNA. Further, a 100-nt-long poly(A) tail was transcribed from the corresponding DNA templates. Uridine 5’-triphosphate (UTP) was replaced with the triphosphate derivative of m1Ψ (m1ΨTP) (TriLink, San Diego, CA, USA Biotechnologies) in the transcription reaction. Capping of the IVT mRNAs was performed co-transcriptionally using the trinucleotide cap1 analog CleanCap (TriLink, San Diego, CA, USA). After DNase digestion, the synthesized mRNA was isolated from the reaction mix by precipitation with half volume of 8 M LiCl solution (Sigma-Aldrich, Hamburg, Germany), and finally the pellet was dissolved in nuclease-free water. The RNA concentration was determined using a Nanodrop 2000c spectrophotometer (Thermo Fisher Scientific). Removal of dsRNA contaminants from 100 to 500 μg IVT mRNA was performed using microcentrifuge spin columns (NucleoSpin Filters, Macherey-Nagel, Düren, Germany), cellulose fibers (C6288, Sigma-Aldrich), and a chromatography buffer containing 10 mM HEPES (pH 7.2), 0.1 mM EDTA, 125 mM NaCl, and 16% (v/v) ethanol according to a previously published protocol^72^.

### Flow cytometric analysis of transfection efficacy

6 hours post-treatment with CART/mRNA polyplexes, NK cells were stained with LIVE/DEAD™ Fixable Yellow Staining Kit (Thermo Fisher Scientific) for 20 minutes at room temperature, then stained with anti-CD3-PE (BioLegend, Clone UCHT1), anti-CD14-APC (BioLegend, Clone HCD14), anti-CD16-PerCP/Cy5.5 (BioLegend, Clone 3G8), and anti-CD56-PE-Cy7 (BioLegend, Clone HCD56) for 20 minutes at room temperature. Data were acquired on an Aurora flow cytometer (Cytek Biosciences) and analyzed by FlowJo version 10.6.1.

### Antibody conjugation, mass cytometry staining, and data acquisition

Antibodies used for mass cytometry staining are shown in Table S1. Antibodies were conjugated using the MaxPar X8 labeling kits (Fluidigm) and lyophilized into single-use pellets for antibody stability prior to use (Biolyph) as previously described^73,74^. NK cells were stained for mass cytometric analysis as previously described^40,59^. Briefly, NK cells were resuspended in cisplatin (Enzo Life Sciences; to stain for cellular viability) for 1 minute and then quenched with 100% FBS. Next, cells from each sample were uniquely barcoded using palladium isotopes as described^75^; 2 out of 4 palladium isotopes were used for each sample, yielding 6 possible combinations. Cells with unique barcodes were then pooled and stained with surface antibodies for 30 minutes at 4°C, fixed (BD FACS Lyse), permeabilized (eBioscience Permeabilization Buffer), and stained with intracellular antibodies for 45 minutes at 4°C. Cells were then suspended in iridium intercalator (Fluidigm) in 2% PFA overnight, followed by 1 wash in PBS, 3 washes in water, and addition of 1X EQ Four Element beads (Fluidigm) immediately before acquisition on a Helios mass cytometer (Fluidigm).

### Mass cytometry data analysis

The open source statistical software R (https://www.r-project.org/) was used for all mass cytometry data analysis. First, bead normalization, bead removal, and de-barcoding were performed using premessa (https://github.com/ParkerICI/premessa). Next, NK cells were gated from debarcoded files according to the scheme in **Supplementary Figure 3**. All data were then transformed using the hyperbolic sine transformation with cofactor equal to 5, to correct for heteroskedasticity. Uniform Manifold Approximation and Projection (UMAP) dimensionality reduction was performed using the package uwot, a native R implementation of UMAP (https://github.com/jlmelville/uwot), using the following parameters: n_neighbors = 10, min_dist = 0.05, spread = 2, metric = cosine. Each variable of input to UMAP was scaled to mean of 0 and variance of 1.

For all heatmap-based visualizations, unweighted pair group method with arithmetic mean was used for agglomerative hierarchical clustering on unscaled data. Median marker expression values were calculated across the analyzed groups and these values were scaled such that the 1^st^ percentile = 0, and the 99^th^ percentile = 1. To perform Louvain clustering, a principal component analysis (PCA) was first performed on unscaled data. The first 6 principal components were then used to calculate the k-nearest neighbors of each cell. Next, this k-nearest neighbors graph was then used to construct a shared nearest neighbors graph by determining the neighborhood overlap between every cell and its 20 nearest neighbors. Next, a smart local moving modularity optimization algorithm^76^ and the Louvain method for community detection^77^ were used with resolution of 0.8 to calculate cluster membership.

The Simpson diversity index was calculated as 1 − *D*, where 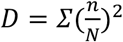, n = the number of cells belonging to cluster i in sample j, and N = the total number of cells in sample j. This index takes into account both the richness and evenness of cluster abundance in each sample.

### *In vitro* NK cell cytokine stimulation and cytotoxicity assays

6 hours post-treatment with CART/mRNA polyplexes, the NK cells were centrifuged, media replaced with complete RP10, and incubated for an additional 12 hours. Then, NK cells were incubated with CellTrace Violet (ThermoFisher Scientific)-stained cancer cell lines K562 or Raji (both cell lines acquired from ATCC, used at passage 2-10, at an effector:target ratio of both 1:3 and 10:1), stimulated with a cocktail of IL-12, IL-15 and IL-18 (rhIL-12, R&D Systems, 5 ng/mL; rhIL-15, Pepro-Tech, 20 ng/mL; rhIL-18, R&D Systems, 50 ng/mL), or left unstimulated without further intervention. 2 hours after addition of cancer cell lines or cytokine stimulation cocktail, brefeldin, monensin (eBioscience), and anti-CD107a-APC-H7 antibody (BD Biosciences, Clone H4A3) were added and the cells incubated for an additional 4 hours. After incubation, cells were stained with LIVE/DEAD™ Fixable Yellow Staining Kit (Thermo Fisher Scientific) for 20 minutes at room temperature, then fixed (2% paraformaldehyde), permeabilized (eBioscience Permeabilization Buffer) and stained with anti-IFNγ-Alexa Fluor 647 (BioLegend, Clone 4S.B3) and anti-TNFα-BV650 (BioLegend, Clone MAb11) for 30 minutes at room temperature. Anti-human CD19 CAR was detected by staining with FITC-conjugated soluble human CD19 (ACROBiosystems) at room temperature for 20 minutes. Data were acquired on an Aurora flow cytometer (Cytek Biosciences) and analyzed by FlowJo version 10.6.1.

## AUTHOR DETAILS

**Aaron J. Wilk**

Department of Medicine, Division of Infectious Diseases and Geographic Medicine; Program in Immunology; Medical Scientist Training Program, Stanford University School of Medicine, Stanford, California

<u>Competing interests:</u> No competing interests declared

<u>Contributed equally with:</u> Nancy L. Benner

**Nancy L. Benner**

Department of Chemistry, Stanford University, Stanford, California

<u>Competing interests:</u> No competing interests declared

<u>Contributed equally with:</u> Aaron J. Wilk

**Rosemary Vergara**

Department of Medicine, Division of Infectious Diseases and Geographic Medicine, Stanford University School of Medicine, Stanford, California

<u>Competing interests:</u> No competing interests declared

**Ole A.W. Haabeth**

Department of Medicine, Division of Oncology, Stanford Cancer Institute, Stanford University, Stanford, California

<u>Competing interests:</u> No competing interests declared

**Ronald Levy**

Department of Medicine, Division of Oncology, Stanford Cancer Institute, Stanford University, Stanford, California

<u>Competing interests:</u> R.L. is a member of the scientific advisory boards for Five Prime, Quadriga, BeiGene, GigaGen, Teneobio, Sutro, Checkmate, Nurix, Dragonfly, Abpro, Apexigen, Spotlight, 47 Inc, XCella, Immunocore, and Walking Fish.

**Robert M. Waymouth**

Department of Chemistry, Stanford University, Stanford, California

<u>Competing interests:</u> No competing interests declared

**Paul A. Wender**

Department of Chemistry and Chemical and Systems Biology, Stanford University, Stanford, California

<u>Competing interests:</u> No competing interests declared

<u>For correspondence:</u>

**Catherine A. Blish**

Department of Medicine, Division of Infectious Diseases and Geographic Medicine, Stanford University School of Medicine, Stanford, California

Chan Zuckerberg Biohub, San Francisco, California

<u>Competing interests:</u> No competing interests declared

<u>For correspondence:</u>

## Author contributions

A.J.W, N.L.B, P.A.W., and C.A.B. developed the concepts and designed the study. A.J.W. and N.L.B. performed the experiments. A.J.W, N.L.B, R.M.W., P.A.W., and C.A.B analyzed data. A.J.W. and N.L.B. prepared the manuscript with input from all authors. A.J.W. performed bioinformatics analyses. R.V. acquired mass cytometry data. O.A.W.H. and R.L. provided reagents.

## ACKNOWLEDGMENTS

This work was supported by the National Cancer Institute (CA031845 to P.A.W. and R35CA19735304 to R.L.) and the National Science Foundation (CHE848280 to P.A.W. and CHE1607092 to R.M.W., organocatalytic polymerization). C.A.B is supported by the Chan Zuckerberg Biohub, NIH/NIDA 5DP1DA04608902, the Burroughs Wellcome Fund Investigators in the Pathogenesis of Infectious Diseases #1016687, and Bill & Melinda Gates Foundation Pilot Grant through the Stanford Human Systems Immunology Center. C.A.B. is the Tashia and John Morgridge Faculty Scholar in Pediatric Translational Medicine from the Stanford Maternal Child Health Research Institute. A.J.W is supported by the Stanford Medical Scientist Training Program (5T32GM007365-44) and the Stanford Bio-X Interdisciplinary Graduate Fellowship. We gratefully acknowledge the lab of Prof. Crystal Mackall for providing cell lines and Dr. Colin McKinlay for materials, including assistance in the synthesis of the mixed-lipid CARTs. We also gratefully acknowledge Dr. Anne-Maud Ferreira and Dr. Tim Blake for materials and helpful discussion.

