## Supplemental Figures 1-6 for "Charge-Altering Releasable Transporters Enable Specific Phenotypic Manipulation of Resting Primary Natural Killer Cells"

**SUPPLEMENTARY INFORMATION**

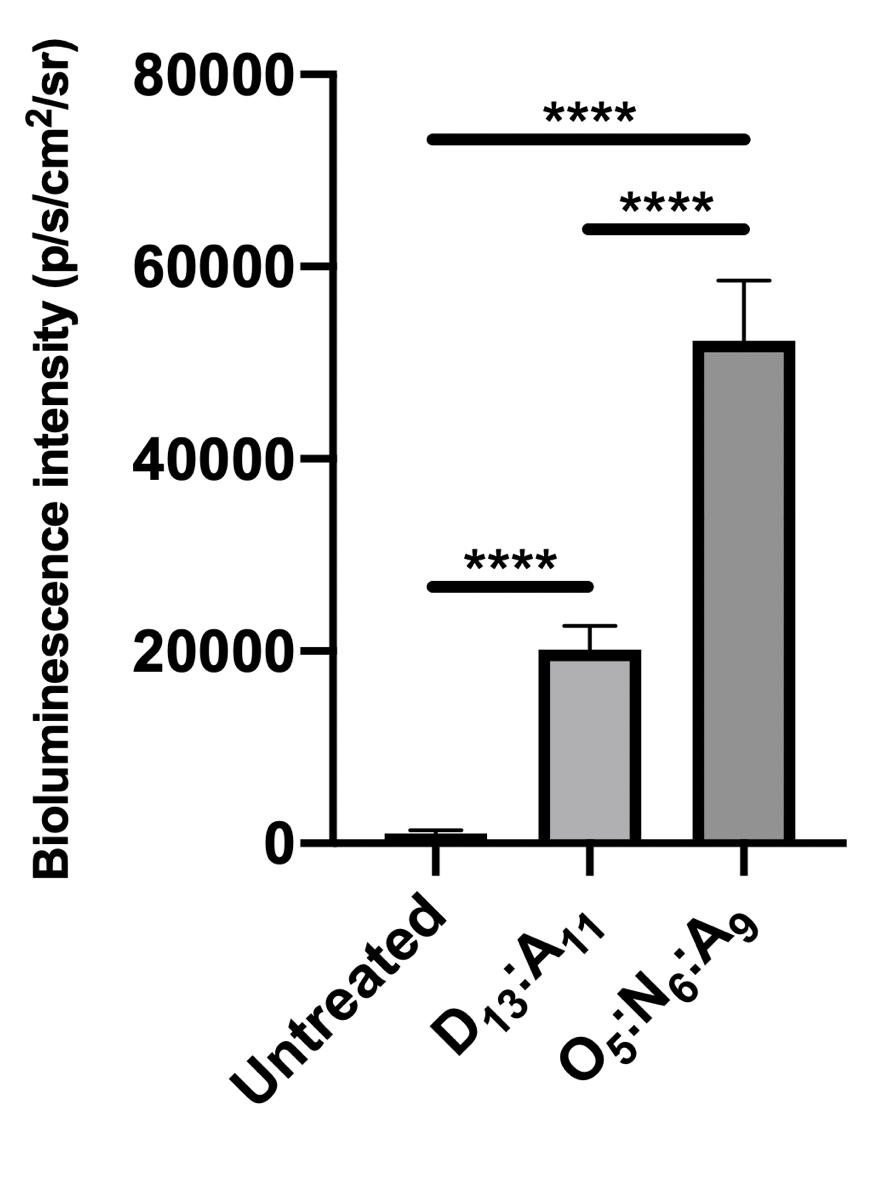

**Supplementary Figure 1. CARTs transfect resting NK cells with luciferase-encoding mRNA**. Quantification of bioluminescence imaging results 6 hours post-transfection of resting primary human NK cells (100,000 cells per well in 100 μL of serum free media) with luciferase-encoding mRNA, n = 4. Error is ± SD. ****, p<0.0001 by unpaired t-test.

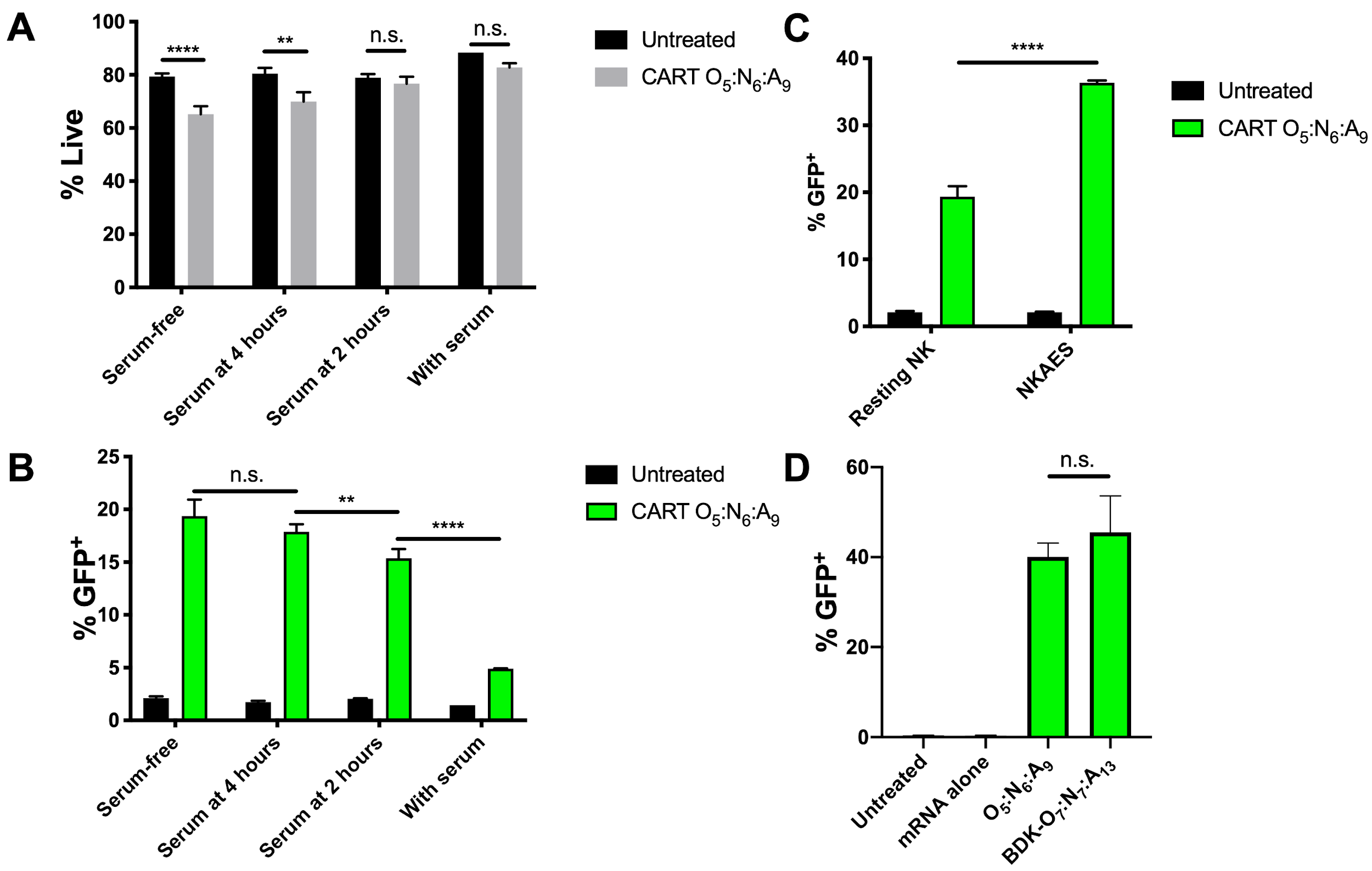

**Supplementary Figure 2. Optimization of CART-mediated transfection of resting primary NK cells. A-B)** Shortening the length of serum-free incubation can improve cell viability without substantially sacrificing transfection efficacy of CART-complexed mRNA. Viability **(A)** and percent transfection **(B)** of primary resting human NK cells (100,000 cells/well in 100 μL serum free RPMI) using CART O_5_:N_6_:A_9_. CART/mRNA polyplexes were formulated at a 10:1 (+/-) charge ratio as previously described. Serum was added to the media at the indicated time points. NK cells were incubated with the complexes for a total of 6 h before analysis by flow cytometry. **C)** Expanded NK cells are more amenable to CART-mediated transfection than resting NK cells. NK cells expanded *via* the NK cell activation and expansion system (NKAES)^56^ as well as resting primary human NK cells from n = 5 donors were transfected with CART O_5_:N_6_:A_9_-complexed GFP mRNA (100,000 cells/well in 100 μL serum free RPMI) and analyzed for GFP expression 6 hours post-transfection. ****, p<0.0001 by paired t-test. **D)** CART O_5_:N_6_:A_9_ or CART BDK-O_7_:N_7_:A_13_ have similar transfection efficacies. Percent transfection of primary resting human NK cells (500,000 cells/well in 100 μL serum free RPMI) using GFP mRNA alone, CART O_5_:N_6_:A_9_ or CART BDK-O_7_:N_7_:A_13_. CART/mRNA polyplexes were formulated at a 10:1 (+/-) charge ratio as previously described. NK cells were incubated with the complexes for 6 h before analysis by flow cytometry. For all panels, error bars represent ± SD. n.s., not significant at p=0.05.

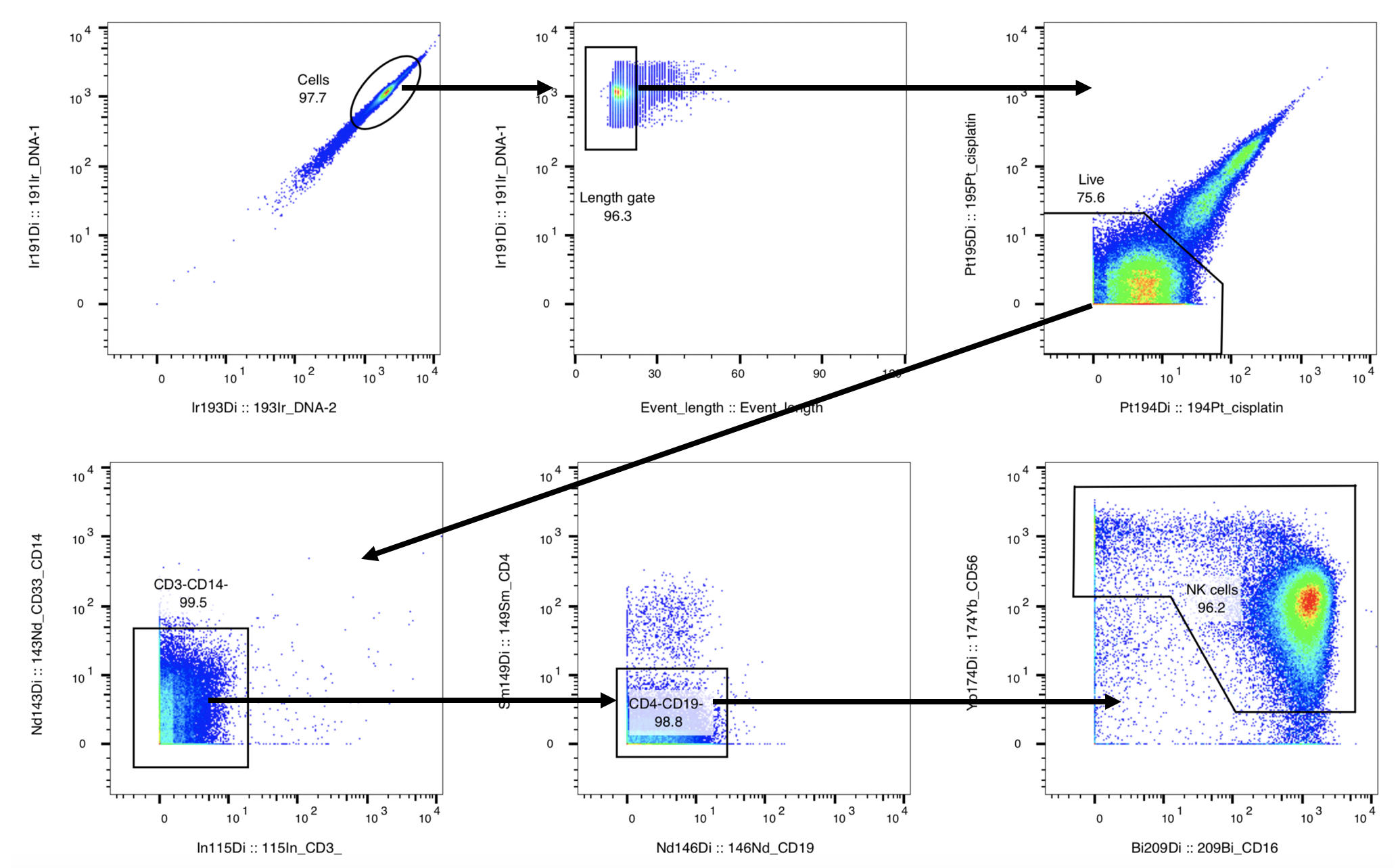

**Supplementary Figure 3. Pre-gating scheme for CyTOF analysis.** Gating scheme used for negative gating of NK cells for mass cytometry analysis. NK cells were enriched by magnetic bead isolation *via* negative selection. Negative gating was used to ensure higher NK cell purity for downstream analysis.

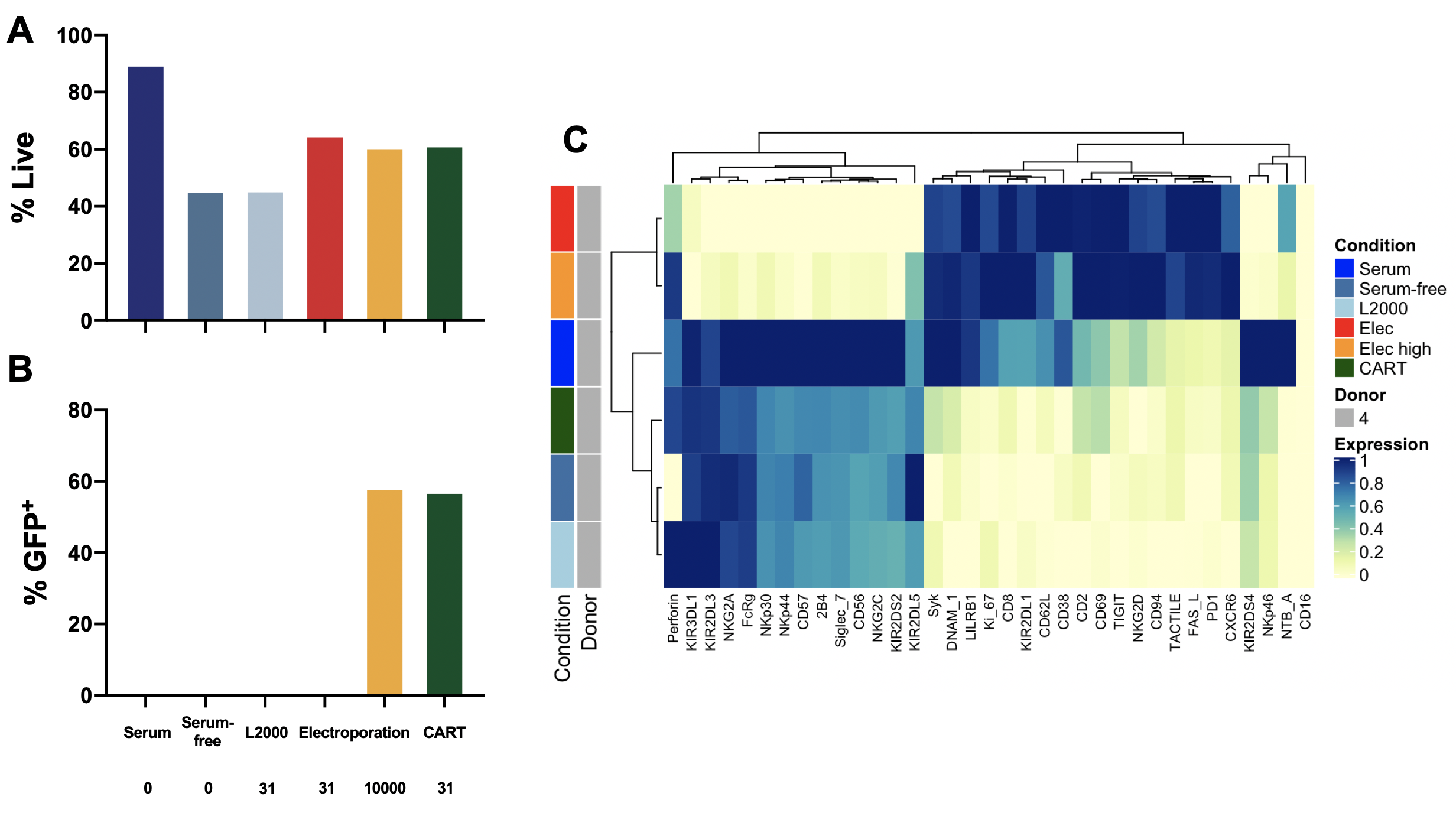

**Supplementary Figure 4. Transfection efficacy, viability, and NK cell proteomic phenotype 6 hours post-transfection with GFP mRNA**. Flow cytometric analysis of NK cell **A)** viability and **B)** transfection efficacy 6 hours post-transfection with GFP-encoding mRNA. **C)** Heatmap of mean marker expression, with samples and markers hierarchically clustered. NK cells were isolated from a single donor.

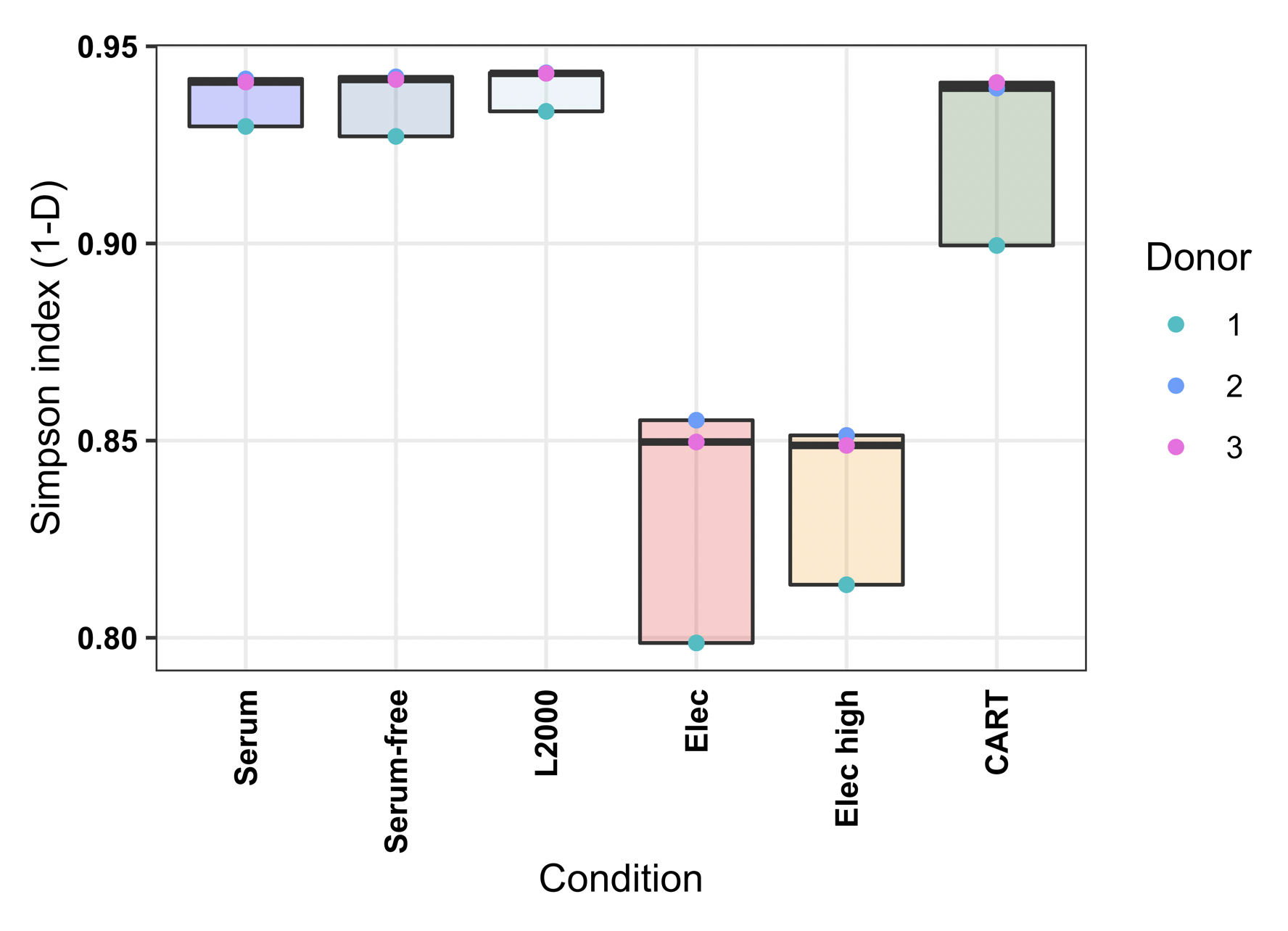

**Supplementary Figure 5. CART transfection, unlike electroporation, maintains comparable NK cell repertoire diversity**. The Simpson diversity index was calculated from mass cytometry cluster abundances 18 hours post-transfection with GFP encoding mRNA.

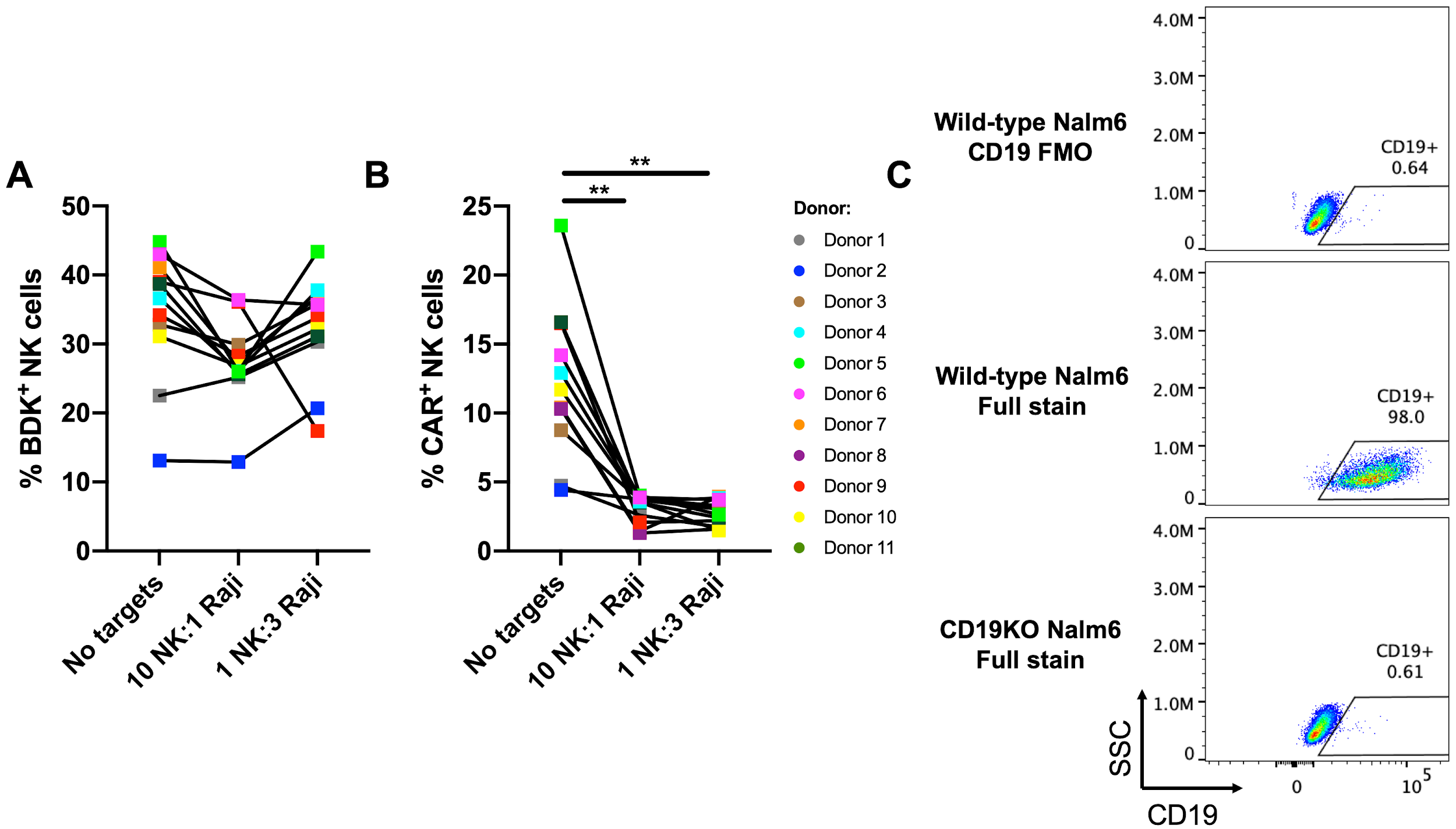

**Supplementary Figure 6. A-B)** Co-culture of anti-CD19 CAR NK cells with CD19^+^ target cells causes internalization of the CAR. **A)** Percentage of BDK^+^ CART-transfected NK cells not co-cultured with Raji target cells, and co-cultured with Raji cells at E:T of 10:1 and 1:3. No significant differences between the groups detected by one-way ANOVA. **B)** Percentage of CAR^+^ CART-transfected NK cells not co-cultured with Raji target cells, and co-cultured with Raji cells at E:T of 10:1 and 1:3. **, p<0.01 by Wilcoxon matched pairs signed-rank test with Bonferroni correction for multiple testing. **C)** Representative flow cytometry plot of wild-type Nalm6 stained without anti-CD19 antibody (CD19 FMO), as well as wild-type and CD19KO Nalm6 stained with the full antibody panel.
